# A ubiquitously expressed UDP-glucosyltransferase, *UGT74J1*, controls basal salicylic acid levels in rice

**DOI:** 10.1101/2021.07.24.453630

**Authors:** Daisuke Tezuka, Hideyuki Matsuura, Wataru Saburi, Haruhide Mori, Ryozo Imai

**Affiliations:** Institute of Agrobiological Sciences, National Agriculture and Food Research Organization (NARO), Kannondai, Tsukuba, 305-8602, Japan; Graduate School of Agriculture, Hokkaido University, Kita-ku, Sapporo 060-8589, Japan; Research Faculty of Agriculture, Hokkaido University, Kita-ku, Sapporo 060-8589, Japan

**Keywords:** Salicylic acid, Glycosylation, UDP-glucosyltransferase, Disease resistance, *Pyricularia oryzae*

## Abstract

Salicylic acid (SA) is a phytohormone that regulates a variety of physiological and developmental processes, including disease resistance. SA is a key signaling component in the immune response of many plant species. However, the mechanism underlying SA-mediated immunity is obscure in rice (*Oryza sativa*). Prior analysis revealed a correlation between basal SA level and blast resistance in a range of rice varieties. This suggested that resistance might be improved by increasing basal SA level. Here, we identified a novel UDP-glucosyltransferase gene, *UGT74J1*, which is expressed ubiquitously throughout plant development. Mutants of *UGT74J1* generated by genome editing accumulated high levels of SA under non-stressed conditions, indicating that UGT74J1 is a key enzyme for SA homeostasis in rice. Microarray analysis revealed that the *ugt74j1* mutants constitutively overexpressed a set of pathogenesis-related (PR) genes. An inoculation assay demonstrated that these mutants had increased resistance against rice blast, but they also exhibited stunted growth phenotypes. To our knowledge, this is the first report of a rice mutant displaying SA overaccumulation.

## 1. Introduction

Salicylic acid (SA), 2-hydroxy benzoic acid, is a small phenolic compound that was first isolated from willow bark. SA functions as a phytohormone and regulates many physiological processes, such as germination, cell growth, respiration, and senescence [1–4]. SA also regulates tolerance against environmental stresses, including heat, chilling, salinity, and oxidative damage [5–8]. However, the most important role of SA is the regulation of disease resistance. The hypersensitive response (HR), an aspect of the plant immune response that is characterized by localized necrosis and tissue lesions [9], is associated with SA accumulation. The HR prevents the penetration and spread of pathogens from the site of infection [10]. Infected local tissues also transmit a signal to distant healthy tissues to induce systemic acquired resistance (SAR) [11,12]. SAR confers resistance to a wide range of pathogens and requires biosynthesis of mobile signals such as methyl salicylate [13].

SA is biosynthesized from chorismic acid via two independent pathways. In the phenylalanine ammonia-lyase (PAL) pathway, cinnamic acid produced by PAL is converted to SA by either β-oxidation or a non-β-oxidation route. In the isochorismic acid synthase (ICS) pathway, isochorismic acid produced by ICS is converted to SA. The intermediate conversion step of the ICS pathway was unknown until a recent study clarified that SA is synthesized from the intermediate isochorismoyl-glutamate in a two-step reaction catalyzed by PBS3 and EPS1 [14]. As an active phytohormone, SA undergoes a variety of modifications, such as methylation, hydroxylation, and sugar conjugation [15]. SA-glycosides are inactive storage forms of SA that accumulate in vacuoles [16–18]. SA 2-*O*-β-glucoside (SAG) and salicyloyl-glucose ester (SGE) have been isolated as major and minor SA glycosides, respectively [19,20].

SA glucosylation is catalyzed by UDP-glucosyltransferases (UGTs). UGT74F1 and UGT76B1 exhibit glucosyltransferase activity toward the hydroxyl group of SA, while UGT74F2 and UGT75B1 show the activity toward its carboxyl group in *Arabidopsis thaliana* [19,21,22]. Tobacco (*Nicotiana tabacum*) SAGT (UGT74G1-like) exhibits transferase activities toward both the hydroxy and carboxyl groups of SA [23], and rice (*Oryza sativa*) OsSGT1 (UGT74H3) catalyzes the glucosylation of the hydroxyl group of SA [24,25]. The UGT74 family proteins function as SA-glucosyltransferases in many plant species, and *UGT74* mutant or overexpressing plants have altered SA and SA glucoside levels. The *ugt74f1* and *ugt74f2 Arabidopsis* single mutants show reduced SAG and SGE levels, respectively, and the *ugt74f1* mutant also shows an increased SA level [19]. Overexpression of *SAGT* in *Nicotiana benthamiana* results in decreased SA and increased SAG levels [26]. The rice *OsSGT1* is a probenazole (PBZ)-inducible gene, and its knock-down mutants exhibit a reduced SAG level under PBZ treatment [25].

The changes in endogenous SA levels before and after pathogen infection differ depending on the plant species. In *Arabidopsis*, tobacco, and cucumber (*Cucumis sativus*), the basal level of SA is low (less than 0.3 µg/g fresh weight (FW)), but its abundance quickly increases in response to pathogen infection (0.5–2.0 µg/g FW) [11,12,27]. In contrast, rice accumulates a relatively large amount of SA under non-stressed conditions (5.0–30.0 µg/g FW), even more than infected *Arabidopsis* or tobacco tissues [28,29]. The SA level in rice does not significantly change following inoculation with either bacterial or fungal pathogens [29]. In addition, a comparative study of rice varieties revealed a positive correlation between basal SA level and general blast resistance [29]. These observations support the hypothesis that the SA level before infection is an important factor that determines rice blast resistance. SA-deficient transgenic *NahG* rice plants exhibit reduced rice blast resistance, further supporting this idea [30]. However, it is not known whether elevated SA levels increase disease resistance in rice, because no rice mutants that constantly overaccumulate SA had been identified.

In this study, we created rice mutants with high basal levels of SA by knocking out a novel UDP-glucosyltransferase gene, *UGT74J1*. We demonstrate that an increased basal SA level contributes to an increase in disease resistance in rice.

## 2. Material and Methods

### 2.1 Phylogenetic analysis

The amino acid sequences of UGTs from *Arabidopsis* and rice were obtained from TAIR (https://www.arabidopsis.org/) and the MSU Rice Genome Annotation Project (http://rice.plantbiology.msu.edu/). The accession numbers of these UGT sequences are listed in Supplemental Table S1. The amino acid sequence alignment was performed using ClustalW (http://clustalw.ddbj.nig.ac.jp/). The phylogenetic tree was drawn by the neighbor-joining method using Geneious software (Biomatters, New Zealand).

### 2.2 Plant materials and growth conditions

Rice cultivar ‘Yukihikari’ was used in this study. The seeds were sterilized with 70% ethanol for 5 min and with 2% sodium hypochlorite for 30 min and subsequently incubated for 2 d in the dark at 30°C for gemination. The germinated seeds were planted in soil and grown in a greenhouse maintained at a 14-h light/10-h dark cycle at 28°C.

The length of the leaf blade and leaf sheath was measured in 50-d-old plants. The heading day was recorded when the first spike emerged from the tip of the leaf sheath.

### 2.3 Semi-quantitative RT-PCR

Plant tissues were sampled at several growth stages. Root, leaf sheath, and leaf blade were collected from 30-d-old plants. Inflorescence and anther were collected from 7- and 9-wk-old plants, respectively. Palea, lemma, and ovary were collected from florets at 5 d after flowering. Total RNA was extracted from each tissue using an RNeasy Plant Mini Kit (Qiagen, Germany), and first-strand cDNA was synthesized from 500 ng total RNA using the PrimeScript RT reagent Kit (Takara, Japan). Semi-quantitative RT-PCR was performed using PrimeSTAR GXL (Takara, Japan). Gene-specific primers are listed in Supplemental Table S2. The following conditions were used for amplification of partial *UGT* fragments: 98°C for 30 s, followed by 30–34 cycles of 98°C for 10 s, 60°C for 15 s, and 68°C for 30 s. The housekeeping gene *OsACT1* was amplified under the following conditions: 98°C for 30 s, followed by 28 cycles of 98°C for 10 s, 55°C for 15 s, and 68°C for 30 s.

### 2.4 Creation of the *ugt74j1* mutants by genome editing

The clustered regularly interspaced short palindromic repeat (CRISPR)/CRISPR-associated protein 9 (Cas9) system was used to create the *ugt74j1* mutants. The CRISPR/Cas9 target sites were located in the first or second exon of *UGT74J1* and were designed using the CRISPRdirect software (https://crispr.dbcls.jp/). The expression vectors for the CRISPR/Cas9 system were constructed as in a previous study [31]. The paired oligonucleotides encoding a 20-nt sequence on the target site were annealed and introduced into pU6gRNA: GTTG<u>GGTTGCTGAGCGTGGAGCTC</u> and AAAC<u>GAGCTCCACGCTCAGCAACC</u> for target site 2; GTTG<u>GTGAGATAAGACCTCAAGCT</u> and AAAC<u>AGCTTGAGGTCTTATCTCAC</u> for target site 3, where the underline indicates the 20-nt target site. Subsequently, the gRNA expression cassettes on pU6gRNA were digested and subcloned into the pZH_gYSA_MMCas9 binary vector using *Pac*I and *Asc*I. Transgenic plants expressing the CRISPR/Cas9 system were generated using an *Agrobacterium tumefaciens*– mediated transformation method [32]. Genomic DNA was extracted from regenerated plants and used for cleaved amplified polymorphic sequence (CAPS) analysis. The amplified fragments derived from the mutants were sequenced using the BigDye Terminatorv3.1 Cycle Sequencing Kit and 3130xl Genetic Analyzer (Thermo Fisher Scientific, USA). Homozygous mutant plants lacking the CRISPR/Cas9 expression cassette were isolated by self-crossing.

### 2.5 Quantification of endogenous SA and SAG

Endogenous SA and SAG levels were determined using the fifth leaf of 25-d-old plants. The leaf tissue was frozen in liquid nitrogen immediately after harvesting and then ground to a fine powder. Sample preparation and measurement using a deuterium-labeled internal control were performed as described previously [33].

### 2.6 Expression analysis using real-time PCR

Total RNA was extracted from the fifth leaf of wild-type and mutant plants using the RNeasy Plant Mini Kit (Qiagen, Germany). First-strand cDNA was synthesized from 500 ng total RNA using the PrimeScript RT reagent kit (Takara, Japan). Quantitative RT-PCR (qRT-PCR) was performed using an ABI7500 Real-Time PCR System (Thermo Fisher Scientific, USA) and TB Green Premix Ex Taq II (Takara, Japan). The following PCR conditions were used: 95°C for 30 s, followed by 40 cycles of 95°C for 5 s and 60°C for 34 s. The transcripts were normalized relative to the transcript levels of the *OsACT1* housekeeping gene. The specificity of the PCR was automatically determined using a melting curve analysis of the amplified products performed by the PCR system. The primers used for the qRT-PCR are listed in Supplemental Table S2.

### 2.7 Detached leaf inoculation assay

The rice cultivar ‘Yukihikari’ carries the *Pia* resistance gene and is susceptible to *Pyricularia oryzae* (race: 007). A spore suspension of *P. oryzae* (race: 007) was prepared as described in a previous study [34]. The fifth leaves of 25-d-old plants were detached and placed on wet paper towels in a plastic box. Each leaf was inoculated with four 3-μL spots of the spore suspension (1.0 × 10^5^ spores/mL) and incubated for 5 d at 25°C. The lesion lengths were measured from digital images of the leaves using ImageJ. The differences in lesion size between the wild type and each mutant were statistically analyzed using a Student’s *t*-test.

## 3. Results

### 3.1 *UGT74J1* is a ubiquitously expressed UDP-glucosyltransferase

To clarify the relationship between basal SA level and disease resistance in rice, we created a mutant that overaccumulates SA. We expected that disrupting SA glucosylation might increase endogenous levels of SA, so we searched the rice genome database, MSU RGAP (http://rice.plantbiology.msu.edu), for a highly and ubiquitously expressed SA-glucosyltransferase gene. Nine unknown UGT74 proteins with significant homology to a previously characterized SA-glucosyltransferase, OsSGT1, were identified in the database (Figure 1). These UGT genes are clustered on chromosomes 4 and 9, suggesting they originated through gene duplications (Supplemental Figure S1).

**Figure 1.**
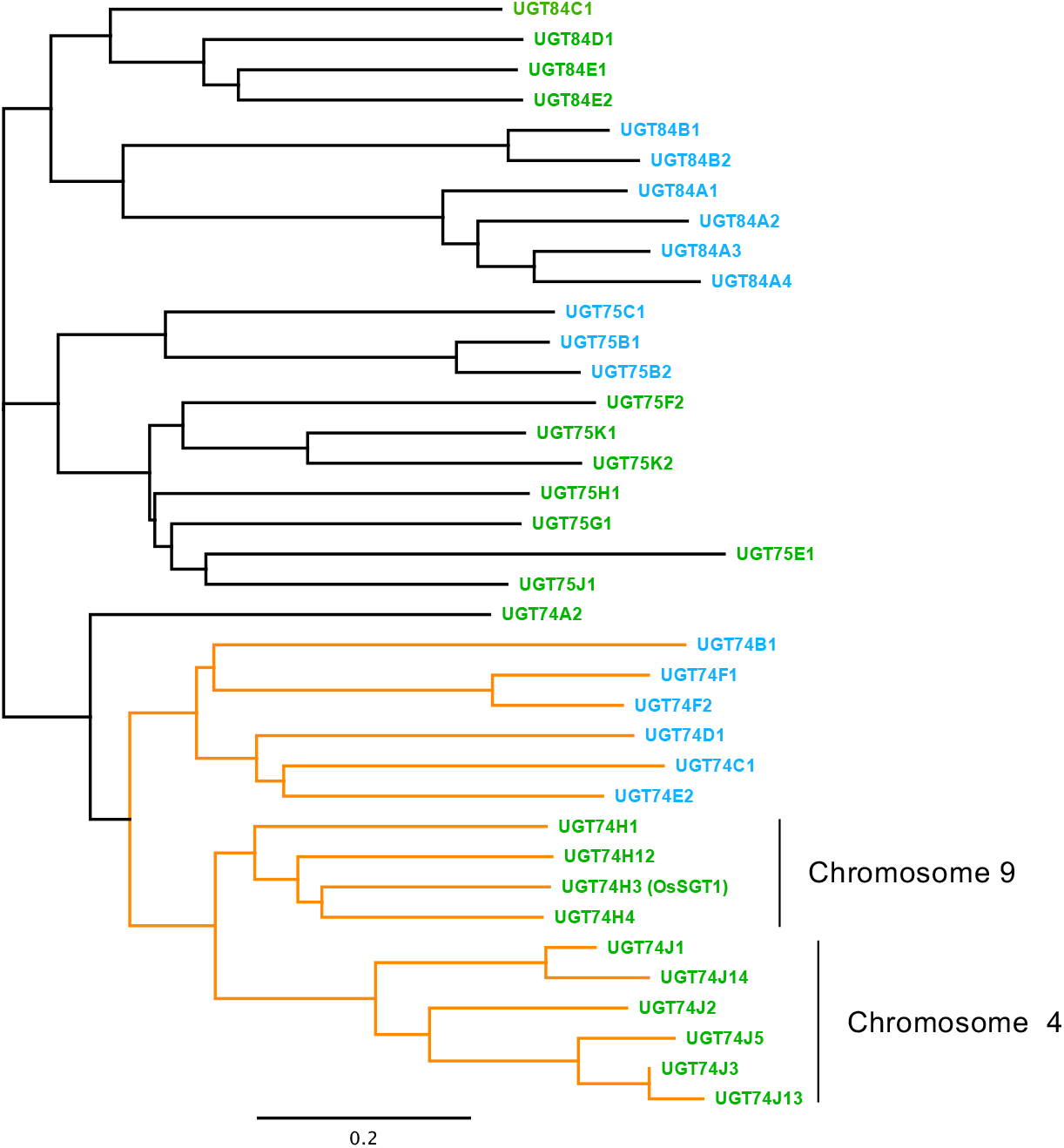
Phylogenetic analysis of UDP-glucosyltransferases in *Arabidopsis* and rice. The amino acid sequences of the UDP-glucosyltransferase family protein domain (accession: PLN02173) were aligned using ClustalW, and a phylogenetic tree was created using the neighbor-joining method. The bar represents the evolutionary distance between the proteins expressed as the number of substitutions per amino acid residue. Green and blue type indicate genes from rice and *Arabidopsis* UGTs, respectively. The clade including known SA-glucosyltransferase (UGT74F1, UGT74F2, and UGT74H3) is indicated by orange lines.

To identify UGT genes that control basal SA levels, we analyzed the tissue-specific expression of the UGT74 family genes under non-stressed conditions (Figure 2). *UGT74J13, UGT74J14, UGT74J2, UGT74J3, UGT74J5*, and *UGT74H12* showed similar patterns of expression, with most mRNA accumulating in the roots, palea, lemma, and ovary. *OsSGT1* (*UGT74H3*) was specifically expressed in the inflorescence and anthers. Expression of *UGT74H1* and *UGT74H4* was observed primarily in leaf blade and in floral organs such as the palea, lemma, and ovary. On the other hand, *UGT74J1* mRNA was detected in all tissues examined, although expression levels were tissue-dependent. We chose *UGT74J1* as a target gene for modulating steady-state SA levels.

**Figure 2.**
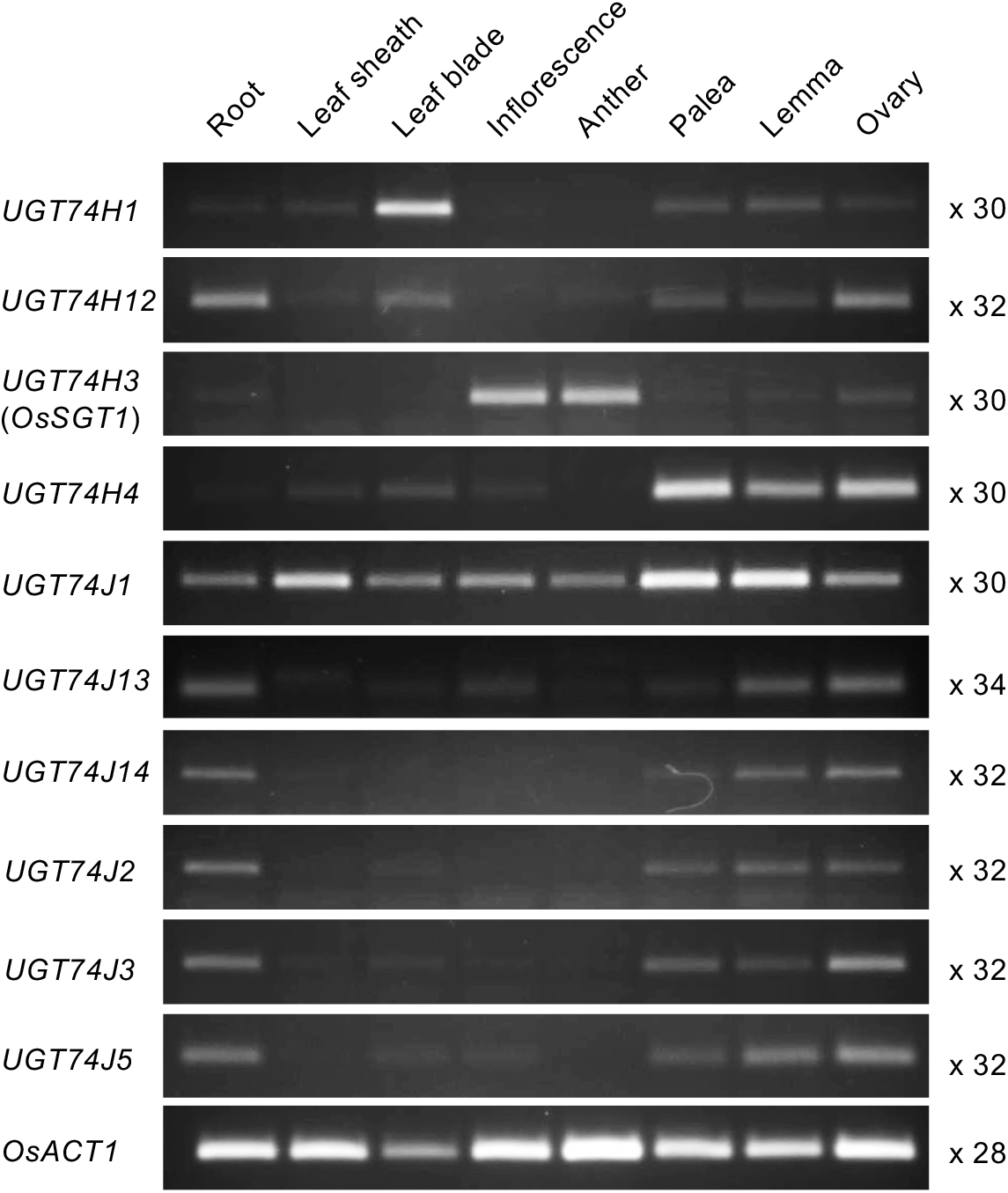
*UGT74j1* is a ubiquitously expressed UDP-glucosyltransferase gene in rice. Each tissue was sampled at the following growth stages: root, leaf sheath, and leaf blade were collected from 30-d-old plants; inflorescence and anther were collected from 7- and 9-wk-old plants, respectively; palea, lemma, and ovary were collected from florets on the fifth day after flowering. Semi-quantitative RT-PCR was performed using gene-specific primers, and the cycle number is indicated at right. *OsACT1* was used as the endogenous control.

### 3.2 Creation of *ugt74j1* knockout mutants using CRISPR/Cas9

*ugt74j1* knockout mutants were created using CRISPR/Cas9-mediated genome editing. Two guide RNA sequences were designed using the CRISPRdirect software (Figure 3A). Transgenic plants expressing Cas9 and either of two targeted guide RNAs were created through *Agrobacterium*-mediated transformation. Genome-edited plants were selected by CAPS and sequencing analysis. We selected two genome-edited T1 plants that had a heterozygous G-nucleotide deletion at target 2 and a heterozygous G-nucleotide insertion at target 3, respectively (Supplemental Figure S2A). Homozygous mutant plants without the transgene integration were selected at the T2 generation (Supplemental Figure S2B,C). Two independent genome edited lines, 2-7 and 3-13, were utilized in the following study (Figure 3B). *UGT74J1* transcript levels were suppressed in both mutants (Figure 3C). This suppression may be caused by nonsense-mediated mRNA decay [35]. Together, these data indicate that lines 2-7 and 3-13 are both knockout mutants of *UGT74J1*.

**Figure 3.**
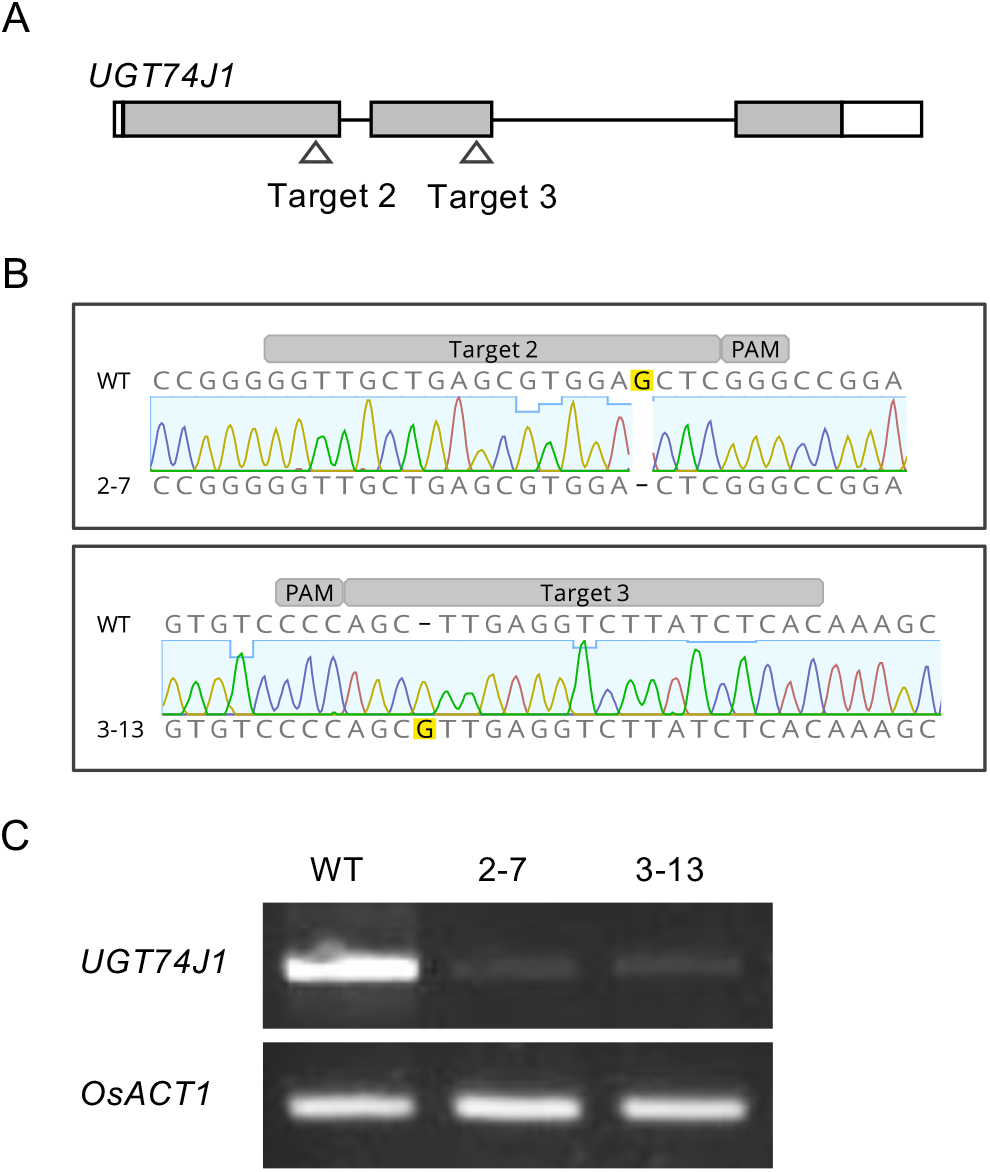
Creation of the *ugt74j1* mutants by genome editing. (A) A schematic model of the CRISPR/Cas9 target sites on UGT74J1. Gray and white boxes indicate the exon and untranslated region of *UGT74J1*. (B) The genotypes of two independent mutants, 2-7 and 3-13. A partial fragment including the CRISPR/Cas9 target site was amplified from the mutants and aligned with the wild-type sequence. (C) The expression of *UGT74J1* in the mutants. Total RNA was extracted from 14-d-old plants. Semi-quantitative RT-PCR was performed using *UGT74J1*-specific primers. *OsACT1* was used as an endogenous control.

### 3.3 *ugt74j1* mutants show SA overaccumulation

We suspected that the *ugt74j1* mutants might have changes in SA level due to the blockage of glucosylation, so we determined SA levels for the mutants by ultra-performance liquid chromatography–tandem mass spectrometry (UPLC-MS/MS). The *ugt74j1* mutants 2-7 and 3-13 exhibited 1.7- and 1.3-fold increases in SA levels, respectively (Figure 4A). We also determined SAG levels in the mutants. Although they showed a tendency towards lower SAG levels, there was no significant difference from that of the wild type (Figure 4B). These data suggest that UGT74J1 regulates basal levels of SA under non-stressed conditions. In addition, the *ugt74j1* mutants showed reduced stature and delayed heading relative to the parental strain under non-stressed conditions (Figure 5A). The leaf sheath and blade were 10–20% shorter than in the wild type (Figure 5B). The heading date of the *ugt74j1* mutants was delayed by 5–6 d, while no significant alterations were observed in flower morphology and fertility (Figure 5C).

**Figure 4.**
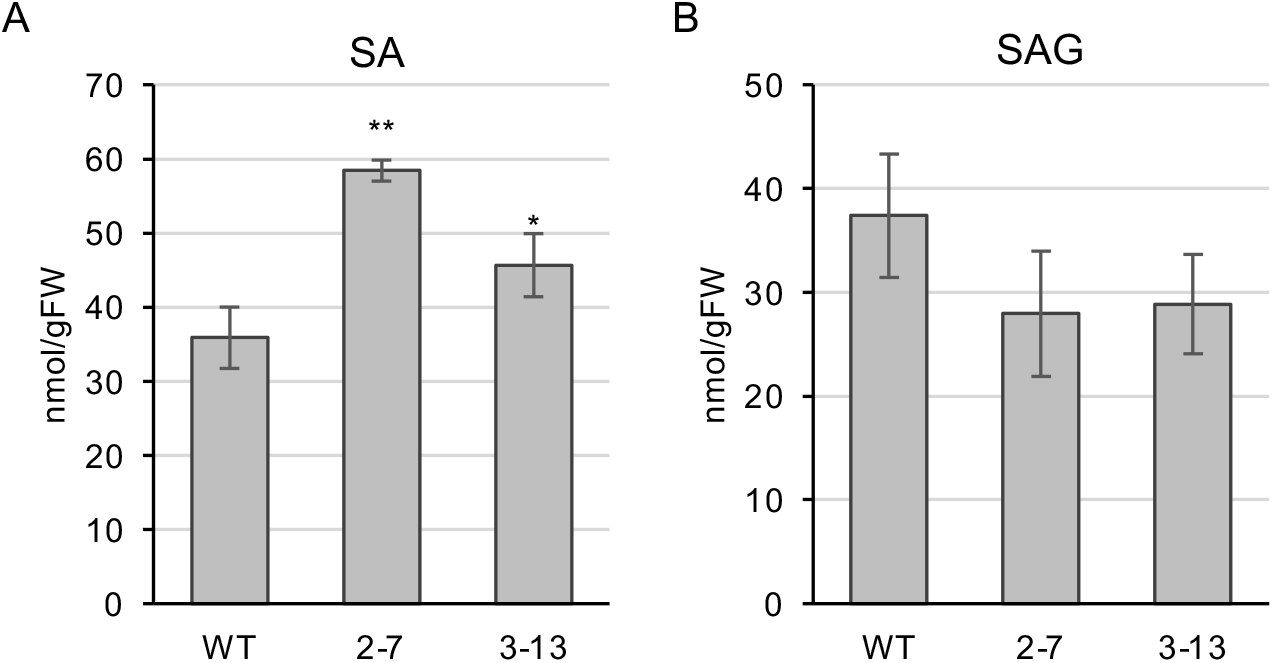
*ugt74j1* mutants overaccumulate SA under non-stressed conditions. (A, B) Salicylic acid (SA) and SA 2-*O*-β-glucoside (SAG) were extracted from the fifth leaf of 25-d-old plants and quantified using UPLC-MS/MS. The data represent the means of four independent experiments ± SE. Significant differences were calculated using Student’s *t*-tests: \**p* < 0.05; \*\**p* < 0.01.

**Figure 5.**
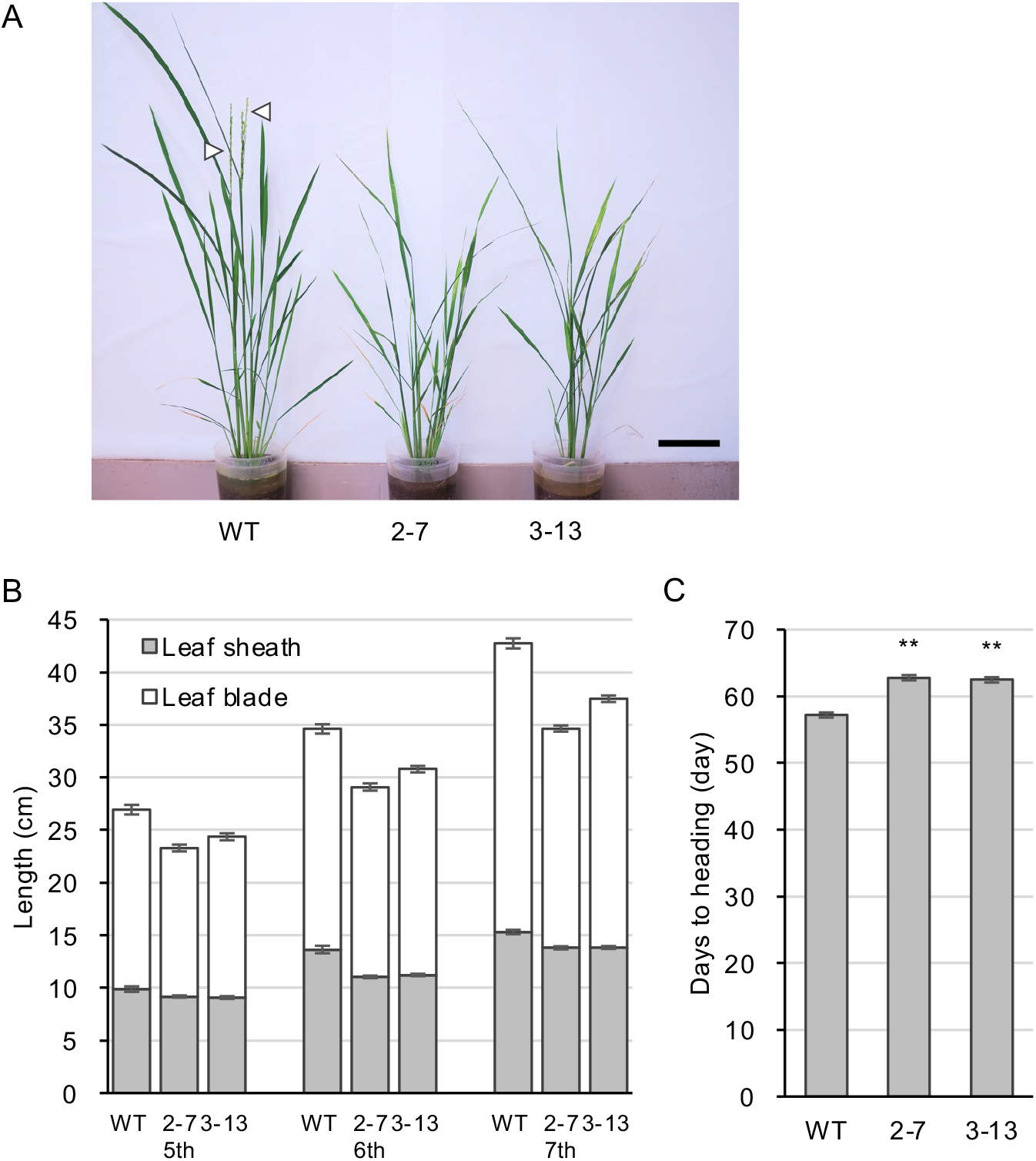
*ugt74j1* mutants show mild stunting and delay in heading. (A) Growth of the *ugt74j1* mutants. Image is of 60-d-old plants. White triangles indicate the headed spikes. The scale bar represents 10 cm. (B) The lengths of the leaf sheath and leaf blade in the mutants. The fifth to seventh leaves of 50-d-old plants were used. The data represent the mean ± SE (*n* = 16). (C) The heading dates of the mutants. The data represent the mean ± SE (*n* = 15). Asterisks indicate statistically significant differences compared with the wild type (*t*-test, \*\**p* < 0.01).

### 3.4 SA overaccumulation upregulates defense-related gene expression

To determine whether the difference in basal SA levels affects gene expression, we compared the transcriptomes of the *ugt74j1* mutant 2-7 and wild-type lines using a microarray. We identified 225 genes whose expression increased more than 5-fold in the mutant (Supplemental Table S3). Among the up-regulated genes with functional annotations, we noticed groups of genes that had specific functions in abiotic stress, hormone biosynthesis, transcription, and defense against pathogens (Table 1). We also observed increased expression of 20 genes encoding pathogenesis-related (PR) proteins, including PR1, PR2, PR3, PR4, PR5, PR8, PR10, and PR13. Moreover, we validated the high expression of several PR genes using real-time PCR in both *ugt74j1* mutants, 2-7 and 3-13, as compared to wild type (Figure 6). There seemed to be a correlation between the basal SA accumulation and the level of PR gene upregulation in these three lines (Figures 4, 6). Together, these data suggested that overaccumulation of SA induces resistance to a broad spectrum of pathogens.

**Table 1.**
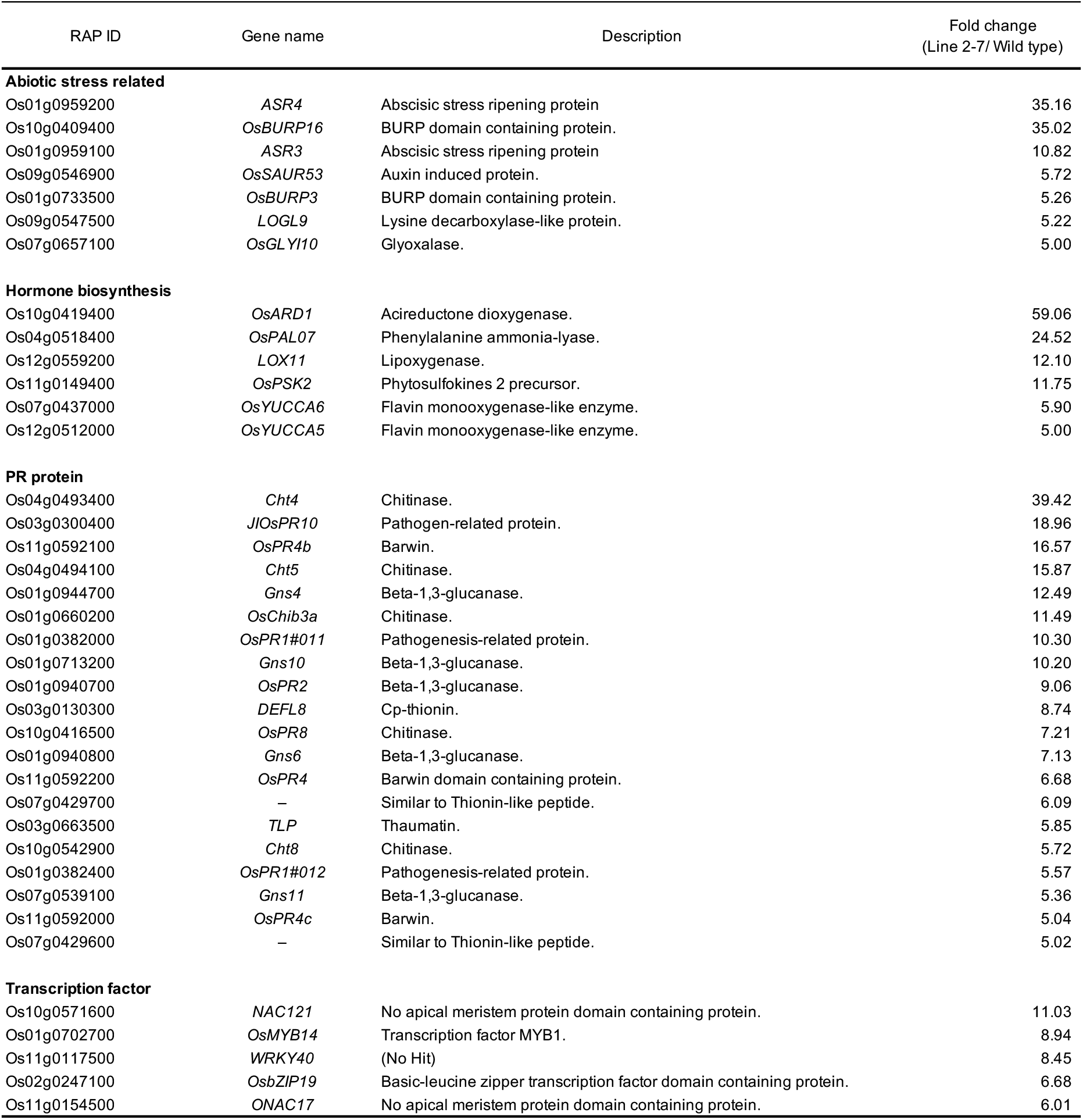
Transcriptome comparison of the *ugt74j1* mutant versus wild type.

**Figure 6.**
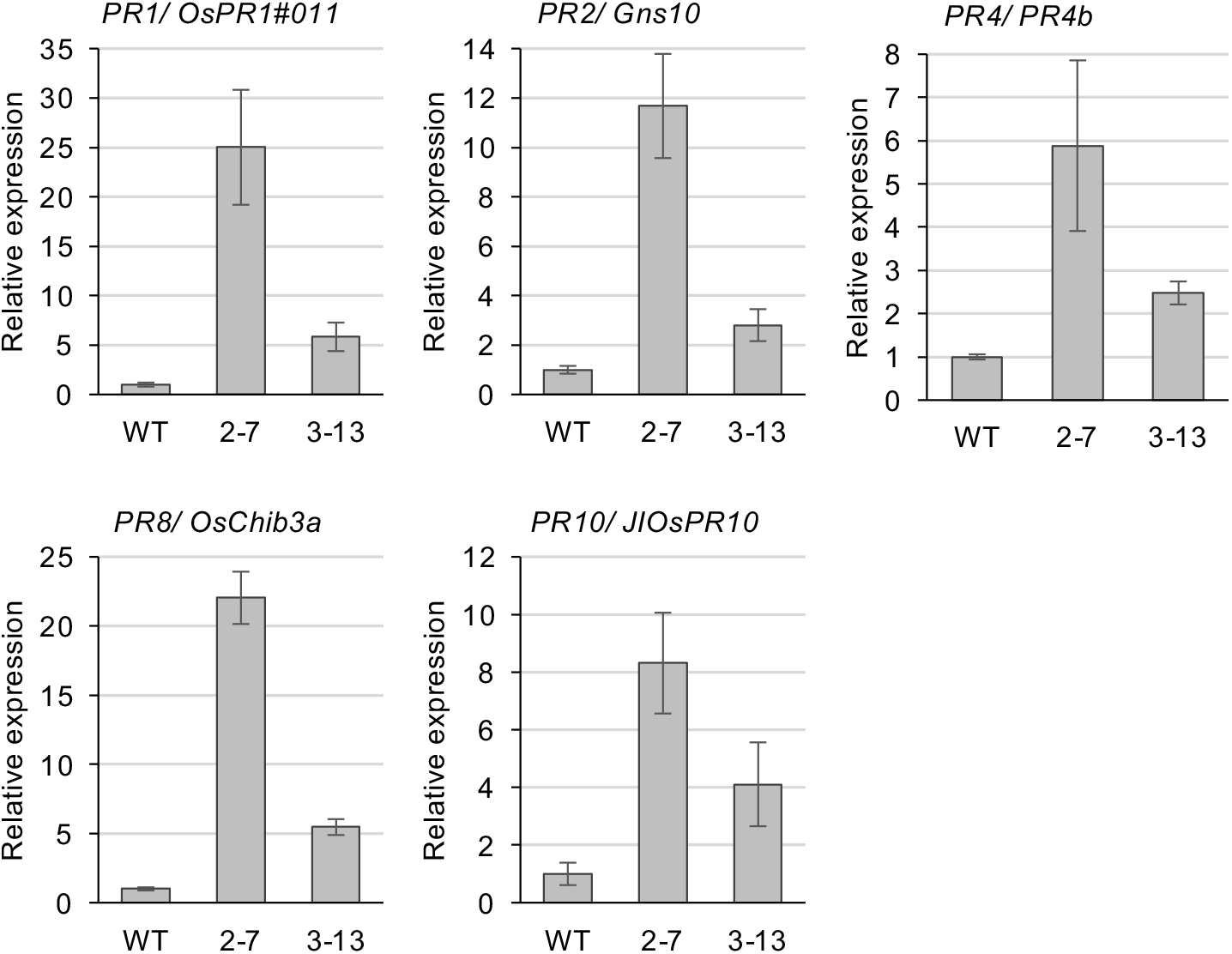
PR gene expression is increased in the *ugt74j1* mutants. qRT-PCR was performed using total RNA isolated from the fifth leaves of the wild type and mutants grown under non-stressed conditions. The expression level of each PR gene was normalized to the expression of *OsACT1*. The data represent the means of four independent experiments ± SE.

### 3.5 *ugt74j1* mutants have increased rice blast resistance

We gauged disease resistance of the *ugt74j1* mutants by a leaf inoculation assay using *P. oryzae*. The detached fifth leaves of the mutants and wild type were inoculated with a spore suspension of *P. oryzae*. After a 5-d incubation, we measured the lesion lengths on the inoculated leaves. The *ugt74j1* mutant lines 2-7 and 3-13 showed a markedly decreased lesion length as compared to wild-type leaves (Figure 7A). The average reduction was 50% and 62%, respectively, and these changes were statistically significant (Figure 7B). Thus, activation of PR genes in response to increased endogenous SA levels likely contributes to enhanced resistance to *P. oryzae*.

**Figure 7.**
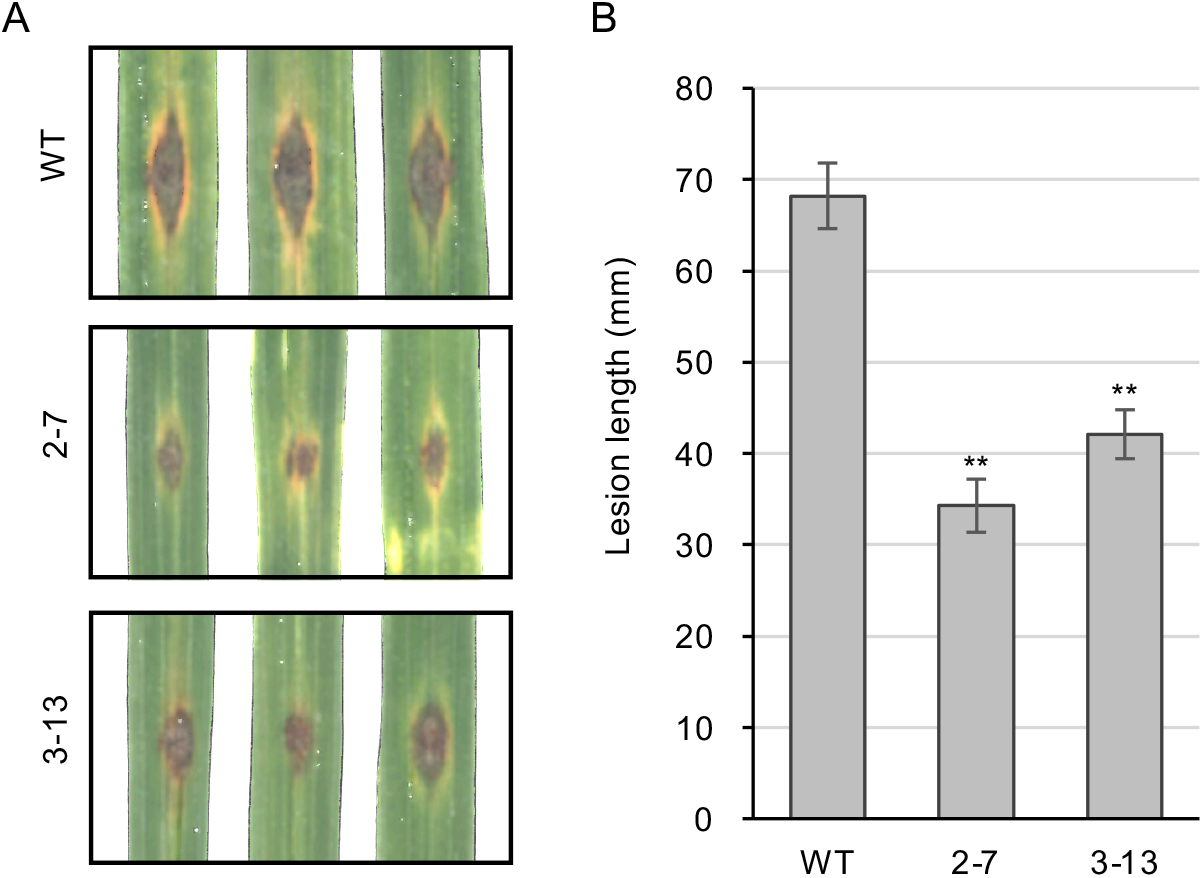
*ugt74j1* mutants have increased resistance to rice blast. (A) Disease resistance assay using *P. oryzae*. The fifth leaves of the wild type and the *ugt74j1* mutants were inoculated with a spore suspension. After 5-d incubation, lesion length was imaged. (B) Quantification of the lengths of the lesions shown in (A). The data are represented as the mean ± SE (*n* = 24). Asterisks indicate statistically significant differences compared with the wild type (*t*-test, *p* < 0.01).

## 4. Discussion

Our expression analysis identified *UGT74J1* as a novel UGT74 group gene that is expressed ubiquitously. In the *ugt74j1* mutants, we observed overaccumulation of SA and a concomitant upregulation of PR genes, and the plants exhibited resistance against rice blast. This supported the previous observation of a positive correlation between basal SA levels and disease resistance in a wide range of rice varieties.

So far, OsSGT1 was the only rice UDP-glucosyltransferase known to have activity toward SA [24,25]. Our data now suggest that UGT74J1 also utilizes SA as a substrate. Our phylogenetic analysis identified ten members in the UDP-glucosyltransferase clade that includes OsSGT1 and UGT74J1 (Figure 1). The sub-clade including OsSGT1 has four members that cluster on chromosome 9, while the other sub-clade, represented by UGT74J1, has six members and clusters on chromosome 4. It is not known whether members of these two sub-clades have distinct enzymatic activities. Among the genes in the chromosome 9 group, OsSGT1 has been functionally characterized as an SA-glucosyltransferase that can transfer a glucose moiety to the hydroxy group of SA [24,25]. In contrast to our *ugt74j1* mutant, *OsSGT1* knockdown plants do not show altered SA levels under non-stressed conditions [25]. This could be explained by the fact that *OsSGT1* is a PBZ-inducible gene [25], and basal expression of *OsSGT1* is maintained at low levels in healthy leaves and roots (Figure 2). Therefore, the contribution of OsSGT1 to the basal SA level appears to be small.

The UGT genes that form a clade with OsSGT1 exhibit tissue-specific expression in leaves (*UGT74H1*), in florets (*UGT74H4*), and in roots and florets (*UGT74J13, UGT74J14, UGT74J2, UGT74J3, UGT74J5, UGT74H12*) (Figure 2).

Therefore, these UGTs are expected to catalyze SA glucosylation in specific tissues. On the other hand, *UGT74J1* was unique in its ubiquitous expression throughout the plant under non-stressed conditions (Figure 2). Since the basal SA levels increased in the *ugt74j1* mutants, we believe that UGT74J1 is a constitutive SA-glucosyltransferase that controls SA–SAG homeostasis under non-stressed conditions (Figure 4).

The endogenous level of SA in the absence of stress is relatively high in rice as compared with other plant species [28,36]. For example, non-stressed leaf tissues contain ∼0.3 µg SA/g FW in *Arabidopsis* [27] and 0.04–0.07 µg SA/g FW in tobacco [38], while in rice (cultivar ‘Nipponbare’) leaves contain 10 µg SA/g FW [25,30]. It is hypothesized that rice has a different mode of action for SA in defense signaling, and a comparative study of basal SA levels in 28 rice varieties revealed that they ranged from 7 µg/g FW to 30 µg/g FW and were positively correlated with blast resistance [29]. This suggests that the blast resistance of a variety could be enhanced by increasing the endogenous SA level. The endogenous SA level of our *ugt74j1* mutant was 7.57 µg/g FW, which was 1.5-fold higher than that of the wild type (cultivar ‘Yukihikari’) at 4.94 µg/g FW (Figure 4). This increase was enough to induce expression of a set of defense-related genes (Table 1) and confer increased resistance against rice blast (Figure 7). The basal SA level of wild-type ‘Yukihikari’ was lower than the range of SA levels observed in the comparative study [29]. This could be explained by differences in the methods utilized for quantifying SA level; we utilized UPLC-MS/MS, which generally allows more specific and accurate measurement than can be obtained using high-performance liquid chromatography with an ultraviolet detector, as in the Silverman study.

The effect of endogenous SA on growth has been studied in *Arabidopsis. Arabidopsis* mutants with high SA levels, such as *cpr6* and *snc1*, exhibit decreased growth in both the roots and the shoots [39,40]. These SA-accumulating mutants contain lower levels of free indole-3-acetic acid (IAA) and have reduced sensitivity to auxin [41]. In contrast to these *Arabidopsis* studies, our transcriptome analysis in rice showed no changes in the expression of many auxin-inducible genes such as *AUX/IAAs* and *ARF*s (data not shown). Additionally, *YUCCA5* and *YUCCA6*, which are involved in IAA synthesis in rice [42], are up-regulated in the *ugt74j1* mutants (Table 1). It is plausible that these genes are induced due to auxin deficiency in the *ugt74j1* mutants, which may explain the stunted growth phenotype of the mutants.

In *Arabidopsis*, SA-mediated immune signaling is almost completely dependent on NPR1 [43]. In contrast, SA signaling in rice is mediated through two independent pathways represented by the key regulators OsNPR1 and WRKY45, respectively [44,45]. When SA or benzothiadiazole (BTH) is exogenously applied, WRKY45 is phosphorylated by a MAPK cascade and converted to an active form [46,47]. The activated WRKY45 auto-regulates its transcription and induces downstream genes such as *WRKY62* and *RDX* [48]. Our microarray analysis did not detect up-regulation of these WRKY genes in the *ugt74j1* mutants (Table 1), suggesting that a WRKY45-mediated SA signal is not activated in the *ugt74j1* mutants. However, we observed up-regulation of some PR genes in the *ugt74j1* mutant (Table 1, Figure 6). One, *OsPR1#11* (*PR-1b*), has been characterized as an OsNPR1-dependent and SA/BTH-inducible gene [44]. Therefore, basal SA accumulation may stimulate SA signaling that is mediated by OsNPR1 but not WRKY45. It is possible that the two SA signaling pathways in rice may differentially regulate SA signals under stressed and non-stressed conditions.

Our study of the *ugt74j* mutants strongly supports the hypothesis that basal SA levels determine disease resistance in rice. Based on this insight, it should be possible to improve the disease resistance of existing rice cultivars by modifying endogenous SA levels. On the other hand, we also showed that excessive SA accumulation has a negative effect on growth. Future studies will need to address how to uncouple these two traits.

## Notes

### Competing Interest Statement

The authors have declared no competing interest.

