## Supplemental Materials for "A ubiquitously expressed UDP-glucosyltransferase, *UGT74J1*, controls basal salicylic acid levels in rice"

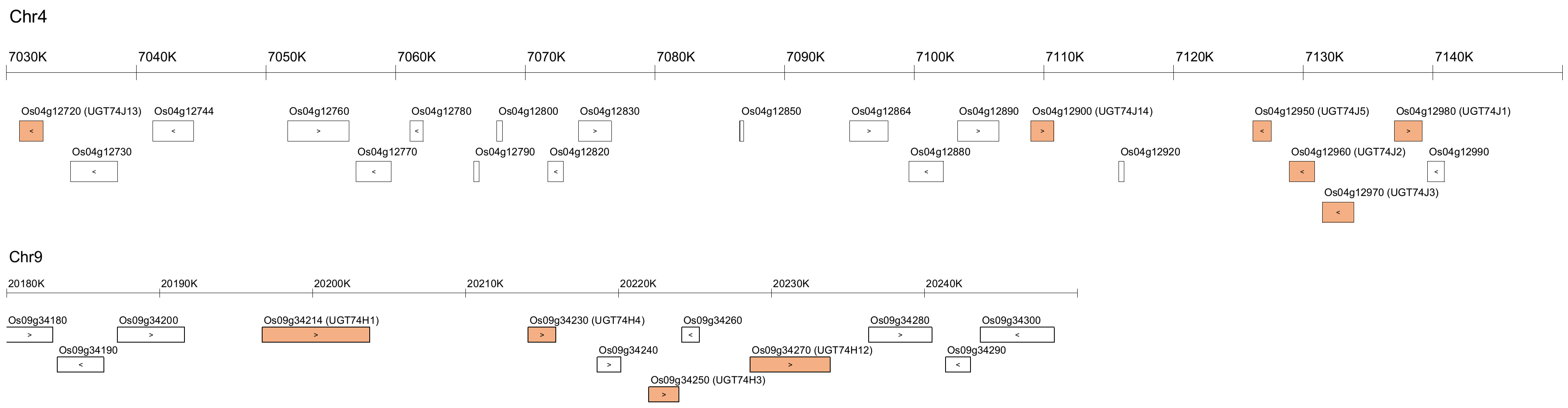


**Figure S1.** Schematic model of the UGT gene clusters on chromosomes 4 and 9.
The UGT genes are indicated by orange boxes. The locus name and chromosomal position of each gene conform to the MSU Rice Genome Annotation Project Database.

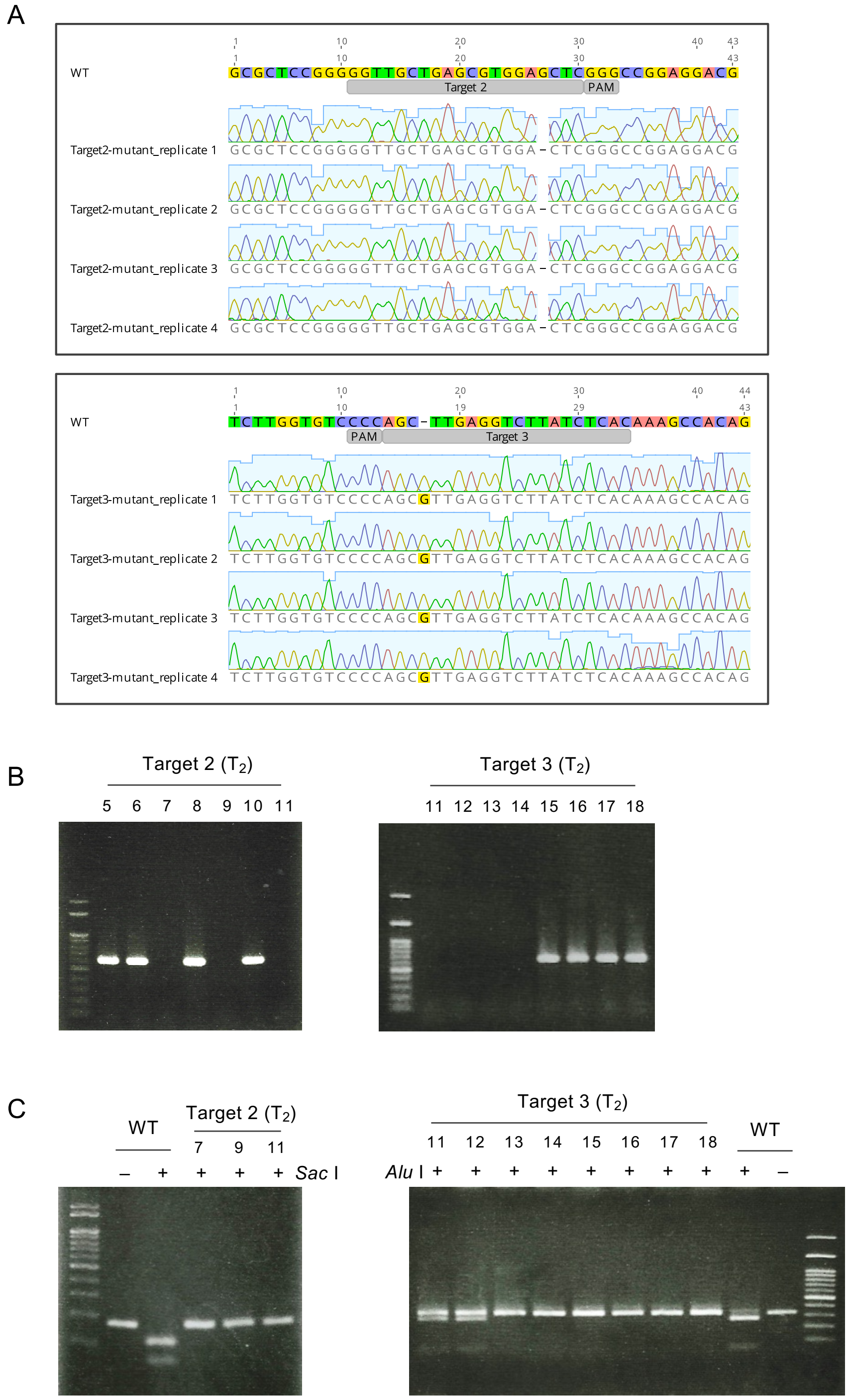


**Figure S2.** Screening for the *ugt74j1* mutants.
(A) Partial sequence of *UGT74J1* in the mutants. Genomic DNA was extracted from two independent $T_1$ plants. The *UGT74J1* fragments were amplified and cloned. Sequences from four independent clones are aligned with the wild type. (B) Segregation of the CRISPR/Cas9 expression cassette in $T_2$ transgenic rice. *Hygromycin phosphotransferase* partial fragments were amplified. (C) Segregation of the *UGT74J1* mutation in $T_2$ transgenic rice. The genotype of each plant was observed by a CAPS assay.

Table S1. Accession numbers of UDP-glucosyltransferases used in phylogenetic analysis

| **Name** | **Species** | **Gene ID** | **Protein ID** |
|---|---|---|---|
| UGT74B1 | *A. thaliana* | At1g24100 | NP_173820.1 |
| UGT74C1 | *A. thaliana* | At2g31790 | NP_180738.1 |
| UGT74D1 | *A. thaliana* | At2g31750 | NP_180734.1 |
| UGT74E2 | *A. thaliana* | At1g05680 | NP_172059.1 |
| UGT74F1 | *A. thaliana* | At2g43840 | NP_973682.1 |
| UGT74F2 | *A. thaliana* | At2g43820 | NP_181910.1 |
| UGT75B1 | *A. thaliana* | At1g05560 | NP_563742.1 |
| UGT75B2 | *A. thaliana* | At1g05530 | NP_172044.1 |
| UGT75C1 | *A. thaliana* | At4g14090 | NP_193146.1 |
| UGT84A1 | *A. thaliana* | At4g15480 | NP_193283.2 |
| UGT84A2 | *A. thaliana* | At3g21560 | NP_188793.1 |
| UGT84A3 | *A. thaliana* | At4g15490 | NP_193284.1 |
| UGT84A4 | *A. thaliana* | At4g15500 | NP_193285.1 |
| UGT84B1 | *A. thaliana* | At2g23260 | NP_179907.1 |
| UGT84B2 | *A. thaliana* | At2g23250 | NP_179906.1 |
| UGT74A2 | *O. sativa* | LOC_Os03g48740 | LOC_Os03g48740.1 |
| UGT74J1 | *O. sativa* | LOC_Os04g12980 | LOC_Os04g12980.1 |
| UGT74J13 | *O. sativa* | LOC_Os04g12720 | LOC_Os04g12720.1 |
| UGT74J14 | *O. sativa* | LOC_Os04g12900 | LOC_Os04g12900.1 |
| UGT74J2 | *O. sativa* | LOC_Os04g12960 | LOC_Os04g12960.1 |
| UGT74J3 | *O. sativa* | LOC_Os04g12970 | LOC_Os04g12970.1 |
| UGT74J5 | *O. sativa* | LOC_Os04g12950 | LOC_Os04g12950.1 |
| UGT74H1 | *O. sativa* | LOC_Os09g34214 | LOC_Os09g34214.1 |
| UGT74H12 | *O. sativa* | LOC_Os09g34270 | LOC_Os09g34270.1 |
| UGT74H3_OsSGT1 | *O. sativa* | LOC_Os09g34250 | LOC_Os09g34250.1 |
| UGT74H4 | *O. sativa* | LOC_Os09g34230 | LOC_Os09g34230.1 |
| UGT75E1 | *O. sativa* | LOC_Os11g04860 | LOC_Os11g04860.1 |
| UGT75F2 | *O. sativa* | LOC_Os11g25990 | LOC_Os11g25990.1 |
| UGT75G1 | *O. sativa* | LOC_Os06g39270 | LOC_Os06g39270.1 |
| UGT75H1 | *O. sativa* | LOC_Os06g39330 | LOC_Os06g39330.1 |
| UGT75J1 | *O. sativa* | LOC_Os02g10880 | LOC_Os02g10880.1 |
| UGT75K1 | *O. sativa* | LOC_Os01g08440 | LOC_Os01g08440.1 |
| UGT75K2 | *O. sativa* | LOC_Os05g08750 | LOC_Os05g08750.1 |
| UGT84C1 | *O. sativa* | LOC_Os02g09510 | LOC_Os02g09510.1 |
| UGT84D1 | *O. sativa* | LOC_Os05g47950 | LOC_Os05g47950.1 |
| UGT84E1 | *O. sativa* | LOC_Os01g49230 | LOC_Os01g49230.1 |
| UGT84E2 | *O. sativa* | LOC_Os01g49240 | LOC_Os01g49240.1 |

Table S2. Primers used in this study

| Gene name | Experiment | RAP ID | MSU ID | Forward primer sequense (5' to 3') | Reverse primer sequense (5' to 3') |
|---|---|---|---|---|---|
| *OsACT1* | Semi-quantitative RT-PCR | Os03g0718100 | LOC_Os03g50890 | TCCATCTTGGCATCTCTCAG | GTACCCTCATCAGGCATCTG |
| *UGT74J1* | Semi-quantitative RT-PCR | Os04g0206700 | LOC_Os04g12980 | GAGTTTGCCTCGAAGTACGC | CTTGTCCAAATATCGTTGAGCA |
| *UGT74J13* | Semi-quantitative RT-PCR | Os04g0204100 | LOC_Os04g12720 | ACAGGAGGAATGCTGCAAGG | TCAGCACCTTGACCTCCATG |
| *UGT74J14* | Semi-quantitative RT-PCR | Os04g0206000 | LOC_Os04g12900 | TGTTAGCGATGCCTCAGTGG | ACTTGATTAGTCATCATCTGGCCA |
| *UGT74J2* | Semi-quantitative RT-PCR | – | LOC_Os04g12950 | GCTCTGTCACGAAGGAGGAG | AACTCCATCTTGTCGTCTGCTA |
| *UGT74J3* | Semi-quantitative RT-PCR | Os04g0206500 | LOC_Os04g12960 | TGGTGACAGGAAGGAGGACT | CAGCACCTTGACCTCCATGA |
| *UGT74J5* | Semi-quantitative RT-PCR | Os04g0206600 | LOC_Os04g12970 | AACTCGACGTTGGAGGCAAT | TCCCGAATGCACCTCTCAAC |
| *UGT74H1* | Semi-quantitative RT-PCR | Os09g0517900 | LOC_Os09g34214 | GGATCGAGGAGGTGATGCG | TGCAGAGAATTTCCCCCACC |
| *UGT74H12* | Semi-quantitative RT-PCR | Os09g0518000 | LOC_Os09g34230 | ATTTCGCGTGCAATAACCGG | GAACTCGTCCTTCCTAGCGG |
| *UGT74H3_OsSGT1* | Semi-quantitative RT-PCR | Os09g0518200 | LOC_Os09g34250 | GGTGTGTGAGGGAGGTGATG | CATCCGACTGTGCCCATTTC |
| *UGT74H4* | Semi-quantitative RT-PCR | Os09g0518400 | LOC_Os09g34270 | GAGGAGGTGGAGAGGAAGGT | TTCCAAATTCATGGCGAGGC |
| *OsACT1* | Real time PCR | Os03g0718100 | LOC_Os03g50890 | TCTCTCTGTATGCCAGTGGTCG | GTCGAGACGAAGGATAGCATGG |
| *OsPR1#011* | Real time PCR | Os01g0382000 | LOC_Os01g28450 | ACGGGCGTACGTACTGGCTA | CTCGGTATGGACCGTGAAG |
| *Gns10* | Real time PCR | Os01g0713200 | LOC_Os01g51570 | CGACGAGAACGGCAAGCCTG | TGGCCTATCACGGGAGCAAC |
| *OsPR4b* | Real time PCR | Os11g0592100 | LOC_Os11g37960 | CGTCTTCTCCAAGATCGACA | CGAACTGGTAGTCGACGATG |
| *OsChib3a* | Real time PCR | Os01g0660200 | LOC_Os01g47070 | TCTACGACGTGCAGAACAACTTCAG | TCCAACTCAACCACTGTGCAAGTAA |
| *JIOsPR10* | Real time PCR | Os03g0300400 | LOC_Os03g18850 | CCTCAGCCATGCCATTCAG | CTTGTCCACGTCCAGGAACTC |

Table S3. The genes whose expression is more than 5-fold upregulated in the mutant.

| RAP ID | Description | Gene name | Function | Fold change (2-7/ wild type) |
|---|---|---|---|---|
| Os03g0115800 | Conserved hypothetical protein. | – | – | 980.51 |
| Os07g0511400 | Hypothetical protein. | – | – | 591.09 |
| Os12g0489300 | Conserved hypothetical protein. | – | – | 438.93 |
| Os10g0118700 | (No Hit) | – | – | 367.60 |
| Os02g0772100 | Conserved hypothetical protein. | – | – | 281.14 |
| Os05g0582200 | Retrotransposon gag protein family protein. | – | – | 227.33 |
| Os03g0377400 | Hypothetical protein. | – | – | 226.96 |
| Os11g0618200 | (No Hit) | – | – | 176.56 |
| Os04g0510000 | Hypothetical protein. | – | – | 161.54 |
| Os01g0692400 | Conserved hypothetical protein. | – | – | 154.33 |
| Os07g0674900 | (No Hit) | – | – | 138.78 |
| Os11g0224300 | Integrase, catalytic region domain containing protein. | – | – | 115.52 |
| Os05g0521800 | BED finger domain containing protein. | – | – | 98.68 |
| Os03g0714800 | En/Spm-like transposon proteins family protein. | – | – | 85.32 |
| Os07g0442800 | Conserved hypothetical protein. | – | – | 79.91 |
| Os08g0316400 | (No Hit) | – | – | 78.65 |
| Os07g0229600 | (No Hit) | – | – | 77.01 |
| Os04g0115200 | Conserved hypothetical protein. | – | – | 76.54 |
| Os01g0845900 | Calcium-binding EF-hand domain containing protein. | – | – | 73.63 |
| Os03g0446000 | (No Hit) | – | – | 70.95 |
| Os04g0575900 | (No Hit) | – | – | 70.51 |
| Os10g0419400 | Submergence induced protein 2. | OsARD1 | Hormone biosynthesis | 59.06 |
| Os12g0539800 | Conserved hypothetical protein. | – | – | 52.82 |
| Os03g0597800 | Non-protein coding transcript, uncharacterized transcript. | – | – | 51.58 |
| Os03g0629800 | Conserved hypothetical protein. | – | – | 46.32 |
| Os06g0565300 | Hypothetical protein. | – | – | 41.21 |
| Os06g0298100 | Conserved hypothetical protein. | – | – | 39.71 |
| Os04g0493400 | Endochitinase A precursor (EC 3.2.1.14) (Seed chitinase A). | cht4 | PR protein | 39.42 |
| Os05g0374700 | Retrotransposon gag protein family protein. | – | – | 38.59 |
| Os07g0137000 | Myb, DNA-binding domain containing protein. | – | – | 38.49 |
| Os10g0524500 | Mandelonitrile lyase-like protein. | OsNP1 | Cell wall associated | 36.61 |
| Os12g0113600 | Hypothetical protein. | – | – | 36.34 |
| Os12g0428300 | Retrotransposon gag protein family protein. | – | – | 35.89 |
| Os01g0959200 | Abscisic stress ripening protein 1. | ASR4 | Abiotic stress related | 35.16 |
| Os10g0409400 | BURP domain containing protein. | OsBURP16 | Abiotic stress related | 35.02 |
| Os07g0598000 | NADPH HC toxin reductase (Fragment). | – | – | 33.00 |
| Os04g0397800 | Non-protein coding transcript, unclassifiable transcript. | – | – | 32.99 |
| Os01g0108500 | (No Hit) | – | – | 31.52 |
| Os04g0249600 | Rhodanese-like domain containing protein. | OsStr6 | Senescence associated | 29.45 |
| Os01g0159200 | Arabinogalactan protein. | OsFLA8 | Cell wall associated | 28.58 |
| Os07g0511100 | Glycine-rich protein precursor. | – | – | 26.80 |
| Os11g0422000 | Non-protein coding transcript, uncharacterized transcript. | – | – | 26.74 |
| Os01g0946600 | Glucan endo-1,3-beta-glucosidase GV (EC 3.2.1.39) ((1->3)-beta-glucan endohydrolase GV) | – | – | 24.90 |
| Os06g0493100 | Hypothetical protein. | OsRALF22 | – | 24.88 |
| Os04g0518400 | Phenylalanine ammonia-lyase 2 (EC 4.3.1.5). | OsPAL07 | Hormone biosynthesis | 24.52 |
| Os12g0117700 | Hypothetical protein. | – | – | 24.30 |
| Os04g0604000 | Actin filament bundling protein P-115-ABP. | VLN4 | Cell development | 23.96 |
| Os03g0688000 | Ribosome-inactivating protein family protein. | – | – | 22.51 |
| Os06g0218600 | Plastocyanin-like domain containing protein. | OsUCL16 | – | 21.39 |
| Os03g0300400 | Pathogen-related protein (JIOsPR10). | JIOsPR10 | PR protein | 18.96 |
| Os12g0247700 | Beta-glucosidase aggregating factor. | OsJAC1 | – | 18.57 |
| Os06g0483200 | Beta-amyrin synthase. | OsOSC6 | – | 17.37 |
| Os06g0267900 | Disease resistance protein family protein. | – | – | 17.37 |
| Os01g0678000 | Conserved hypothetical protein. | – | – | 17.29 |
| Os11g0576400 | BED finger domain containing protein. | Os_F0247 | – | 17.11 |
| Os01g0946500 | Glucan endo-1,3-beta-glucosidase GV (EC 3.2.1.39) ((1->3)-beta-glucan endohydrolase GV) | – | – | 17.06 |
| Os08g0155900 | (No Hit) | OsDR10 | Biotic stress related | 16.70 |
| Os11g0592100 | Barwin. | OsPR4b | PR protein | 16.57 |
| Os08g0355600 | Hypothetical protein. | – | – | 16.56 |
| Os04g0397900 | (No Hit) | – | – | 16.43 |
| Os05g0409500 | MtN21 protein. | OsUMAMIT9 | – | 16.36 |
| Os04g0494100 | Endochitinase A precursor (EC 3.2.1.14) (Seed chitinase A). | cht5 | PR protein | 15.87 |
| Os02g0129000 | Conserved hypothetical protein. | – | – | 15.25 |
| Os02g0513700 | Protein of unknown function DUF659 domain containing protein. | – | – | 15.08 |
| Os11g0260000 | Hypothetical protein. | – | – | 15.01 |
| Os10g0343500 | (No Hit) | – | – | 14.80 |
| Os02g0576400 | Conserved hypothetical protein. | OsEnS-39 | – | 14.60 |
| Os05g0135500 | Haem peroxidase, plant/fungal/bacterial family protein. | prx71 | – | 14.32 |
| Os08g0367300 | Conserved hypothetical protein. | – | – | 14.08 |
| Os04g0677300 | Harpin-induced 1 domain containing protein. | – | – | 13.87 |
| Os06g0658400 | (No Hit) | – | – | 13.80 |
| Os01g0111700 | Conserved hypothetical protein. | – | – | 13.08 |
| Os06g0591200 | Conserved hypothetical protein. | – | – | 13.04 |
| Os03g0346200 | Conserved hypothetical protein. | – | – | 12.78 |
| Os01g0944700 | Glucan endo-1,3-beta-glucosidase GII precursor (EC 3.2.1.39) ((1->3)- beta-glucan endohydrolase GII) | Gns4 | PR protein | 12.49 |
| Os04g0326000 | ARM repeat fold domain containing protein. | – | – | 12.39 |
| Os03g0251000 | Plant lipid transfer/seed storage/trypsin-alpha amylase inhibitor domain containing protein. | – | – | 12.31 |
| Os08g0277200 | Cinnamoyl-CoA reductase (EC 1.2.1.44). | – | – | 12.30 |
| Os12g0559200 | Lipoxygenase (EC 1.13.11.12). | LOX11 | Hormone biosynthesis | 12.10 |
| Os04g0295500 | Conserved hypothetical protein. | – | – | 11.83 |
| Os11g0149400 | Phytosulfokines 2 precursor | OsPSK2 | Hormone biosynthesis | 11.75 |
| Os12g0508400 | Hypothetical protein. | – | – | 11.71 |
| Os01g0660200 | Acidic class III chitinase OsChib3a precursor (Chitinase) (EC 3.2.1.14). | OsChib3a | PR protein | 11.49 |
| Os06g0602400 | ATP-dependent RNA helicase-like protein. | OsRH52A | – | 11.32 |
| Os11g0632200 | Conserved hypothetical protein. | – | – | 11.24 |
| Os08g0124500 | Resistance protein candidate (Fragment). | – | – | 11.17 |
| Os10g0571600 | No apical meristem (NAM) protein domain containing protein. | NAC121 | Transcription factor | 11.03 |
| Os03g0115700 | Short-chain dehydrogenase/reductase SDR family protein. | – | – | 10.97 |
| Os12g0170800 | 24 kDa protein SC24 (24 kDa seed coat protein). | – | – | 10.84 |
| Os01g0959100 | Abscisic stress ripening protein 1. | ASR3 | Abiotic stress related | 10.82 |
| Os07g0417800 | Galactosyl transferase family protein. | – | – | 10.82 |
| Os01g0382000 | Pathogenesis-related protein 1 precursor (PR-1). | OsPR1#011 | PR protein | 10.30 |
| Os01g0713200 | Beta-1,3-glucanase precursor. | Gns10 | PR protein | 10.20 |
| Os06g0265100 | Hypothetical protein. | – | – | 10.08 |
| Os04g0227500 | DSBA oxidoreductase family protein. | – | – | 10.03 |
| Os07g0599600 | Hypothetical protein. | – | – | 9.49 |
| Os04g0204000 | Conserved hypothetical protein. | – | – | 9.34 |
| Os12g0548700 | CI2C. | – | – | 9.22 |
| Os01g0940700 | Beta-1,3-glucanase (Fragment). | OsPR2 | PR protein | 9.06 |
| Os01g0702700 | Transcription factor MYB1. | OsMYB14 | Transcription factor | 8.94 |
| Os03g0130300 | Cp-thionin. | DEFL8 | PR protein | 8.74 |
| Os04g0509000 | Protein kinase domain containing protein. | – | – | 8.73 |
| Os11g0117500 | (No Hit) | WRKY40 | Transcription factor | 8.45 |
| Os04g0259800 | Conserved hypothetical protein. | – | – | 8.40 |
| Os07g0457200 | Non-protein coding transcript, putative npRNA. | – | – | 8.25 |
| Os02g0602300 | Conserved hypothetical protein. | – | – | 8.21 |
| Os01g0779800 | Hypothetical protein. | – | – | 8.12 |
| Os05g0124000 | Ankyrin repeat containing protein. | OsSTA138 | – | 8.06 |
| Os04g0422300 | Protein of unknown function DUF6 domain containing protein. | – | – | 8.05 |
| Os01g0160900 | Leucine-rich repeat, plant specific containing protein. | – | – | 8.04 |
| Os03g0102700 | Beta-expansin precursor. | OsEXPB7 | Cell wall associated | 7.96 |
| Os01g0863100 | Conserved hypothetical protein. | – | – | 7.95 |

| | | | | |
|---|---|---|---|---|
| Os03g0600600 | Beta-1,3-glucanase precursor. | – | – | 7.95 |
| Os02g0288400 | MRP-like ABC transporter. | OsMRP5 | – | 7.94 |
| Os11g0514000 | Hypothetical protein. | – | – | 7.93 |
| Os07g0677500 | Peroxidase POC1. | POX3006 | | 7.92 |
| Os05g0263300 | Non-protein coding transcript, unclassifiable transcript. | – | – | 7.90 |
| Os02g0561800 | (No Hit) | – | – | 7.79 |
| Os09g0268600 | Zn-finger, PMZ type domain containing protein. | – | – | 7.70 |
| Os06g0261300 | Hypothetical protein. | – | – | 7.66 |
| Os03g0838400 | Ammonium transporter. | AMT3;2 | – | 7.53 |
| Os12g0637400 | Purple acid phosphatase (EC 3.1.3.2) (Fragment). | – | – | 7.51 |
| Os11g0227700 | RPR1. | – | – | 7.32 |
| Os06g0142300 | Early nodulin 93 ENOD93 protein family protein. | OsENOD93 | – | 7.29 |
| Os04g0368000 | (No Hit) | OsWAK43 | – | 7.28 |
| Os10g0416500 | Chitinase 1 precursor (EC 3.2.1.14) (Tulip bulb chitinase-1) (TBC-1). | OsPR8 | PR protein | 7.21 |
| Os05g0550300 | Nonspecific lipid transfer protein. | OsLTP2.4 | – | 7.18 |
| Os05g0164400 | (No Hit) | – | – | 7.18 |
| Os01g0940800 | Beta-1,3-glucanase precursor. | Gns6 | PR protein | 7.13 |
| Os02g0561000 | (No Hit) | – | – | 7.13 |
| Os04g0340100 | Protein kinase-like domain containing protein. | – | – | 7.09 |
| Os01g0510200 | Conserved hypothetical protein. | – | – | 7.05 |
| Os04g0543600 | Amino acid/polyamine transporter I family protein. | OsCAT5 | – | 7.05 |
| Os02g0232900 | Major intrinsic protein. | OsNIP1;1 | – | 7.02 |
| Os04g0648000 | Non-protein coding transcript, uncharacterized transcript. | – | – | 6.79 |
| Os11g0513500 | Hypothetical protein. | – | – | 6.75 |
| Os10g0415100 | Amino acid/polyamine transporter II family protein. | OsGAT4 | – | 6.71 |
| Os09g0424200 | Carbamoyl-phosphate synthase, GATase region domain containing protein. | – | – | 6.68 |
| Os02g0247100 | Basic-leucine zipper (bZIP) transcription factor domain containing protein. | OsbZIP19 | Transcription factor | 6.68 |
| Os11g0592200 | Barwin domain containing protein. | OsPR4 | PR protein | 6.68 |
| Os10g0569400 | RIR1a protein precursor. | Rir1b | Biotic stress related | 6.59 |
| Os02g0560600 | (No Hit) | – | – | 6.49 |
| Os10g0567900 | F-box protein interaction domain containing protein. | – | – | 6.44 |
| Os02g0778900 | Proteinase inhibitor I9, subtilisin propeptide domain containing protein. | – | – | 6.44 |
| Os10g0527400 | Glutathione S-transferase TSI-1 (EC 2.5.1.18) (Glutathione S- transferase 1). | OsGSTU19 | – | 6.42 |
| Os03g0151500 | Hypothetical protein. | – | – | 6.27 |
| Os04g0168300 | Hypothetical protein. | – | – | 6.26 |
| Os01g0701700 | SAM dependent carboxyl methyltransferase family protein. | – | – | 6.24 |
| Os10g0467800 | Cellulose synthase. | OsCesA7 | Cell wall associated | 6.18 |
| Os08g0459400 | Transposase, IS4 domain containing protein. | – | – | 6.10 |
| Os07g0429700 | Similar to Thionin-like peptide. | – | – | 6.09 |
| Os02g0699000 | TGF-beta receptor, type I/II extracellular region family protein. | OsNPF7.2 | – | 6.04 |
| Os11g0154500 | No apical meristem (NAM) protein domain containing protein. | ONAC17 | Transcription factor | 6.01 |
| Os01g0577600 | Protein kinase domain containing protein. | – | – | 6.00 |
| Os10g0570200 | RIR1b protein precursor. | – | – | 5.98 |
| Os04g0105200 | Conserved hypothetical protein. | – | – | 5.96 |
| Os10g0188000 | Cyclin-like F-box domain containing protein. | OsFbox543 | – | 5.96 |
| Os01g0127600 | Bowman-Birk type proteinase inhibitor D-II precursor (IV). | – | – | 5.95 |
| Os07g0437000 | Flavin monoxygenase-like protein floozy. | OsYUCCA6 | Hormone biosynthesis | 5.90 |
| Os03g0165400 | Relative to SR12 protein (Fragment). | OsBgal1 | – | 5.86 |
| Os03g0663500 | Thaumatin, pathogenesis-related family protein. | TLP | PR protein | 5.85 |
| Os02g0561400 | (No Hit) | – | – | 5.83 |
| Os04g0468600 | Conserved hypothetical protein. | – | – | 5.83 |
| Os10g0141600 | Conserved hypothetical protein. | OsFbox531 | – | 5.81 |
| Os06g0592500 | Ethylene-responsive transcriptional coactivator. | – | – | 5.78 |
| Os02g0740900 | WD40-like domain containing protein. | OsWD40-54 | – | 5.76 |
| Os07g0622400 | Conserved hypothetical protein. | – | – | 5.74 |
| Os09g0546900 | Auxin induced protein. | OsSAUR53 | Abiotic stress related | 5.72 |
| Os10g0542900 | Chitinase (EC 3.2.1.14) (Fragment). | Cht8 | PR protein | 5.72 |
| Os10g0555900 | Beta-expansin precursor. | OsEXPB3 | Cell wall associated | 5.71 |
| Os12g0491400 | (No Hit) | – | – | 5.67 |
| Os02g0209900 | (No Hit) | – | – | 5.62 |
| Os11g0282700 | Homeodomain-like containing protein. | – | – | 5.61 |
| Os01g0382400 | Pathogenesis-related protein PRB1-2 precursor. | OsPR1#012 | PR protein | 5.57 |
| Os08g0467300 | ANTH domain containing protein. | – | – | 5.55 |
| Os08g0507800 | (No Hit) | – | – | 5.55 |
| Os01g0722800 | Dimethylmenaquinone methyltransferase family protein. | – | – | 5.54 |
| Os11g0513000 | (No Hit) | – | – | 5.52 |
| Os01g0392600 | Hypothetical protein. | – | – | 5.51 |
| Os01g0822900 | Lipid transfer protein. | OsLTP1.2 | – | 5.50 |
| Os11g0677500 | Retrotransposon gag protein family protein. | – | – | 5.48 |
| Os05g0214300 | MtN3 and saliva related transmembrane protein family protein. | OsSWEET3a | – | 5.48 |
| Os11g0676500 | NBS-LRR type resistance protein (Fragment). | – | – | 5.46 |
| Os01g0310700 | Hypothetical protein. | – | – | 5.46 |
| Os01g0613800 | Peptidase C1A, papain family protein. | – | – | 5.44 |
| Os04g0561500 | Prolyl endopeptidase (EC 3.4.21.26) (Post-proline cleaving enzyme) (PE). | OsPOP9 | – | 5.41 |
| Os03g0179700 | Protein of unknown function DUF567 family protein. | – | – | 5.36 |
| Os07g0539100 | Glycoside hydrolase, family 17 protein. | Gns11 | PR protein | 5.36 |
| Os04g0200700 | Conserved hypothetical protein. | – | – | 5.33 |
| Os04g0121800 | Non-protein coding transcript, uncharacterized transcript. | – | – | 5.32 |
| Os01g0567200 | Conserved hypothetical protein. | – | – | 5.30 |
| Os12g0638100 | Receptor-like protein kinase. | – | – | 5.30 |
| Os04g0408600 | Protein of unknown function DUF662 family protein. | – | – | 5.28 |
| Os11g0513900 | Conserved hypothetical protein. | – | – | 5.27 |
| Os01g0944500 | Glycoside hydrolase, family 17 protein. | – | – | 5.27 |
| Os01g0281600 | Plastocyanin-like domain containing protein. | OsENODL4 | – | 5.26 |
| Os01g0733500 | Dehydration-responsive protein RD22 precursor. | OsBURP3 | Abiotic stress related | 5.26 |
| Os03g0449000 | (No Hit) | – | – | 5.25 |
| Os09g0499400 | Hypothetical protein. | – | – | 5.23 |
| Os04g0677000 | Conserved hypothetical protein. | – | – | 5.22 |
| Os01g0105300 | BED finger domain containing protein. | – | – | 5.22 |
| Os09g0547500 | Lysine decarboxylase-like protein. | LOGL9 | Abiotic stress related | 5.22 |
| Os03g0196400 | Protein of unknown function DUF588 family protein. | – | – | 5.20 |
| Os06g0271000 | UDP-glucuronosyl/UDP-glucosyltransferase family protein. | – | – | 5.19 |
| Os04g0619800 | Conserved hypothetical protein. | – | – | 5.18 |
| Os12g0543800 | (No Hit) | – | – | 5.16 |
| Os07g0263300 | Hypothetical protein. | – | – | 5.15 |
| Os01g0823700 | Plant protein of unknown function DUF641 domain containing protein. | – | – | 5.14 |
| Os09g0538900 | Conserved hypothetical protein. | – | – | 5.12 |
| Os06g0582600 | Cysteine proteinase. | – | – | 5.10 |
| Os09g0491100 | Beta-primeverosidase (EC 3.2.1.149). | OsBGlu30 | – | 5.10 |
| Os04g0309600 | Sucrose synthase. | SUS5 | – | 5.09 |
| Os05g0563600 | Beta-Ig-H3/fasciclin domain containing protein. | OsFLA6 | Cell wall associated | 5.09 |
| Os09g0129500 | Conserved hypothetical protein. | – | – | 5.08 |
| Os01g0611000 | Protein of unknown function DUF642 family protein. | – | – | 5.06 |
| Os04g0627900 | Translation initiation factor SUI1 family protein. | UNP1 | – | 5.05 |
| Os06g0277000 | (No Hit) | – | – | 5.05 |
| Os11g0600900 | Mannose-6-phosphate isomerase (ManA). | – | – | 5.05 |
| Os03g0342100 | Hypothetical protein. | – | – | 5.05 |
| Os11g0592000 | Barwin. | OsPR4c | PR protein | 5.04 |
| Os06g0146600 | Hypothetical protein. | – | – | 5.03 |
| Os07g0429600 | Similar to Thionin-like peptide. | – | PR protein | 5.02 |
| Os11g0125900 | Nucleoside phosphatase GDA1/CD39 family protein. | – | – | 5.01 |
| Os07g0657100 | Glyoxalase/bleomycin resistance protein/dioxygenase domain containing protein. | OsGLYI10 | Abiotic stress related | 5.00 |
| Os12g0512000 | Flavin monoxygenase-like protein floozy. | OsYUCCA5 | Hormone biosynthesis | 5.00 |
